# Population genomics, resistance, pathogenic potential, and mobile genetic elements of carbapenem-resistant *Klebsiella pneumoniae* causing infections in Chile

**DOI:** 10.1101/2022.11.28.517876

**Authors:** Marcelo Veloso, Joaquín Acosta, Patricio Arros, Camilo Berríos-Pastén, Roberto Rojas, Macarena Varas, Miguel L. Allende, Francisco P. Chávez, Pamela Araya, Juan Carlos Hormazábal, Rosalba Lagos, Andrés E. Marcoleta

## Abstract

Multidrug and carbapenem-resistant *K. pneumoniae* (CR-*Kp*) are considered critical threats to global health and key traffickers of resistance genes to other pathogens. In Chile, although a sustained increase in CR-*Kp* infections has been observed, few strains have been described at the genomic level, lacking molecular details of their resistance and virulence determinants and the mobile elements mediating their dissemination. In this work, we studied the antimicrobial resistance and performed a comparative genomics analysis of ten CR-*Kp* isolates from the Chilean surveillance of carbapenem-resistant *Enterobacteriaceae*. High resistance to most of the antibiotics tested was observed among the isolates, five ST25, three ST11, one ST45, and one ST505, which harbored a total of 44 plasmids, many of them predicted to be conjugative and carrying genes conferring resistance to a variety of antibiotic, metals, and disinfectants. Ten plasmids encoding either KPC-2, NDM-1, or NDM-7 carbapenemases were characterized, including novel plasmids with increased resistance gene load and a novel genetic environment for *bla*_KPC-2_ gene. This corresponds to the first report of ST25 and ST45 *Kp* producing NDM-7 in South America, and of an ST505 CR-*Kp* worldwide, producing both NDM-7 and KPC-2. Moreover, we characterized a variety of genomic islands carrying virulence and fitness factors. These results provide baseline knowledge for the detailed understanding of molecular and genetic determinants behind antibiotic resistance and virulence of *K. pneumoniae* in Chile and South America.

## BACKGROUND

Multidrug-resistant (MDR) and hypervirulent *Klebsiella pneumoniae*, particularly carbapenem-resistant strains (CR-*Kp*) causing high mortality and morbidity, are critical concerns[1]. These high-risk pathogens have developed mainly by acquiring different mobile genetic elements (MGEs), including plasmids and genomic islands (GIs) encoding an array of virulence and antibiotic resistance factors[2].

Carbapenems are last-resort antibiotics for treating severe infections caused by MDR *Enterobacteriaceae*[3]. The most concerning carbapenem resistance mechanism corresponds to carbapenem-inactivating beta-lactamases, especially the *K. pneumoniae* carbapenemase (KPC) distributed worldwide, showing the highest prevalence. Moreover, this pathogen is a host and a key trafficker of various other carbapenemases, including NDM, VIM, and OXA-48, all encoded in MGEs[1,2,4].

In Chile, after identifying the first introduced KPC-producing *K. pneumoniae* infection in 2012, the Chilean Public Health Institute (ISP) established clinical surveillance of carbapenem resistance and carbapenemases in *Enterobacteriaceae*. Fifty-eight percent of the 489 carbapenemase-producing isolates processed between 2014 and 2017 corresponded to *Klebsiella* spp., while the predominant carbapenemases corresponded to KPC (77%), NDM (15%), and VIM (7%)[5]. Furthermore, between 2012 and 2020, a sustained increase in resistance to meropenem, ertapenem and imipenem was observed in *K. pneumoniae[6]*.

Despite the high incidence of CR-*Kp* infections in Chile, little is known about these strains’ genomic and molecular features. Moreover, we lack detailed information regarding the phylogenetic relationships among Chilean CR-*Kp* and the remaining South American and global *K. pneumoniae* population, the virulence and resistance factors hosted by these strains, and the mobile elements mediating their dissemination. Furthermore, current public genome databases, including NCBI, lack complete genome sequences of CR-*Kp* isolated in Chile, hampering their inclusion in large-scale population genomic studies to place them in a global epidemiological context.

This work characterized ten CR-*Kp* strains from the Chilean CR *Enterobacteriaceae* surveillance, determining their phenotypic antibiotic resistance, complete genome sequence, virulence and resistance gene profile, and mobile genetic elements associated with these factors. Our results provide baseline knowledge of the virulence and resistance determinants hosted by *K. pneumoniae* strains causing infections in Chile, and the mobile genetic platforms that could mediate their dissemination in our regional context.

## METHODS

### Isolate collection and antimicrobial susceptibility tests

The ten CR-*Kp* isolates described here were collected in different hospitals from two regions of Chile in 2018 and 2019 (Supplementary Table 1). Antimicrobial susceptibility testing was performed by broth microdilution and epsilometry, following the M100 Performance Standards for Antimicrobial Susceptibility Testing, 31st edition. Tigecycline resistance was interpreted according to EUCAST 2019 guidelines. The Blue Carba, boronic acid, and Hodge tests were made following standard procedures.

### Genome sequencing

Genomic DNA was extracted using the GeneJET Kit (Thermo Scientific) and quantified using a Qubit fluorimeter (Invitrogen). Illumina sequencing (100-bp paired-end) was performed by Macrogen Inc. (Korea) using the TruSeq Nano DNA kit and a Hiseq4000 machine. FastQC v0.11.9 [7] and Trimmomatic v0.36 [8] were used for quality check and trimming. Nanopore libraries prepared with the Native Barcoding Expansion (EXP-NBD104) and the 1D Ligation sequencing (SQK-LSK109) kits were run on a MinION device and FLO-MIN106 (R9.4) flow cells. After base-calling with Guppy v5.0.7, the raw Nanopore reads were corrected, trimmed, and posteriorly assembled using Canu v2.3 [9]. Hybrid assemblies were performed using Unicycler v0.4.9b [10], using as input the trimmed short and long reads plus the long reads assembly made with Canu. The assemblies were evaluated using QUAST v5.0.2 [11] and CheckM[12], and annotated with RAST v2.0 [13].

### Phylogenomic analyses

A curated database including 3,443 *K. pneumoniae* sensu stricto genomes from the NCBI assembly database was constructed. Species and sequence types were assigned using Kleborate[14]. Whole-genome-based distance trees were inferred using Mashtree v1.2.0 [15], combining 1000 bootstrap iterations into a final Newick file using RAxML v8.2.12[16] (GTRCAT model). For cgMLST-based phylogenetic analysis, we used Roary [17] to generate a concatenated alignment of the core protein sequences (≥95% identity cutoff) and RaxML to infer a maximum likelihood distance tree (GAMMAPROT model, 1000 bootstrap iterations).

### Identification of resistance and virulence genes

ARG prediction was made by combining RGI v5.2.0 (CARD database v3.1.4) [18] and AMRFinderPlus v3.10.18 [19], harmonizing and consolidating both outputs using hAMRonization v1.0.3 (https://github.com/pha4ge/hAMRonization), following ARG classification according to the CARD ontology. Kleborate ARG prediction was also considered. Metal and biocide resistance gene identification was performed using Diamond blastX [20] and the BACMET database (experimentally confirmed) v2.0 [21]. Virulence factors were predicted using VFanalyzer and the VFDB Database [22]. Additionally, Kleborate was used to search for characteristic *K. pneumoniae* virulence factors.

### Mobile genetic elements analyses

Plasmid identification and typing were made using PlasmidVerify [23] and PlasmidFinder v2.0.1 [24]. Transfer origin (oriT) and conjugation genes detection was performed using oriTfinder [25]. Integrons and insertion sequences were predicted using IntegronFinder v1.5.1 [26] and ISfinder [27]. tDNA classification and prediction of integrated MGEs was performed using our tool Kintun-VLI (https://github.com/GMI-Lab/Kintun-VLI).

Prophage identification was made using PHASTER [28]. MGE clustering to define non-redundant GIs and prophages was performed with CD-HIT [29] (≥80% identity and ?85% coverage).

## RESULTS

### Phenotypic AMR f CR-*Kp* causing infections in Chile

We performed an in-depth characterization of ten CR-*Kp* isolates collected in 2018 and 2019 as part of the ISP surveillance for CR *Enterobacteriaceae* from patients of varying ages, gender, and different infected tissues (Supplementary Table 1). MIC determinations indicated that all of them were resistant to 3^rd^ and 4^th^ generation cephalosporins (most also to combinations with beta-lactamase inhibitors), as well as to meropenem and ertapenem, while seven were resistant to imipenem (Table 1). PCR analysis showed either NDM or KPC carbapenemase genes in six strains, while two encoded both, and two lacked these genes (Supplementary Table 2). Furthermore, Blue Carba, Triton Hodge, and Boronic acid tests confirmed carbapenemase production in the positive isolates, while ESBL production was detected in all of them. Additionally, most isolates were resistant to ciprofloxacin, sulfamethoxazole, and gentamicin, while only VA833 showed resistance to amikacin, two to tigecycline, and two to colistin. Thus, the Chilean isolates corresponded to extensively drug-resistant *K. pneumoniae* with high resistance to carbapenems and other last-resort antimicrobial chemotherapy.

**Table 1.** Minimum inhibitory concentrations for different antibiotics.

| Isolate | FEP | CAZ | CRO | CZA | C/T | TZP | ETP | IPM | MEM | TGC | CIP | AMK | GEN | SXT | CST |
| --- | --- | --- | --- | --- | --- | --- | --- | --- | --- | --- | --- | --- | --- | --- | --- |
| VA4 | >16* | >16* | >4* | ND | ND | >256/4* | >1* | >8* | >32* | 1* | >2* | <=4 | >8* | >=4/76* | <=1 |
| VA32 | >16* | >16* | >4* | 2/4 | >32/4* | >256/4* | >1* | >8* | >32* | 0.25 | >2* | <=4 | <=0,5 | >=4/76* | <=1 |
| VA126 | >256* | >256* | >32* | ND | ND | >256/4* | 16* | 0.25 | 4* | 0.5 | >2* | <=4 | <=0,5 | >=4/76* | <=2 |
| VA172 | >16* | >16* | >32* | 1/4 | >32/4* | >256/4* | 4* | 2* | 2* | 0.5 | >2* | <=4 | <=0,5 | >=4/76* | 4* |
| VA564 | >16* | >16* | >32* | 4/4 | >32/4* | >256/4* | 4* | 0.5 | 4* | <=0,25 | >2* | <=4 | >8* | >=4/76* | <=0,25 |
| VA569 | >16* | >16* | >32* | 16/4* | >32/4* | >256/4* | >32* | >8* | >8* | <=0,25 | >2* | <=4 | >8* | >=4/76* | <=0,25 |
| VA591 | >16* | >16* | >32* | ND | ND | 128/4* | 32* | >8* | 8* | 0.5 | 2* | <=4 | >8* | <=1/19 | >4* |
| VA681 | >16* | >16* | >32* | <=0,5/4 | >32/4* | 128/4* | 8* | <=1 | 4* | 0.5 | >2* | <=4 | >8* | >=4/76* | <=0,25 |
| VA684 | >16* | >16* | >32* | >16/4* | >32/4* | 64/4* | 4* | 8* | 8* | 0.5 | <=0,064 | <=4 | <=0,5 | >=4/76* | <=0,25 |
| VA833 | >16* | >16* | >32* | >16/4* | >32/4* | >256/4* | >32* | >8* | >8* | 2* | >2* | >32* | >8* | >=4/76* | <=0,25 |

\*: Resistant. FEP: cefepime; CAZ: ceftazidime; CRO: ceftriaxone; CZA: ceftazidime/avibactam; C/T: Ceftolozane/Tazobactam; TZP: Piperacillin/Tazobactam; ETP: ertapenem; IPM: imipenem; MEM: meropenem; TGC: tigecycline; CIP: ciprofloxacin; AMK: amikacin; GEN: gentamicin; SXT: sulfamethoxazole; CST: colistin.

### Genomic features of Chilean CR-*Kp* and relationships with *K. pneumoniae* isolated worldwide

We determined the complete genome of each isolate by combining Illumina and Nanopore data, obtaining a closed chromosome and two to six plasmids (Table 2). According to MLST analysis using Kleborate, all belonged to the *K. pneumoniae sensu stricto* (Kp1) species, five from ST25, three ST11, one ST45, and one ST505. Accordingly, a phylogenomic analysis including 3443 Kp1 genomes from the NCBI database (Supplementary Table 3) using Mashtree, showed the Chilean isolates clustering inside four deep-branched groups following the ST (Figure 1A). The three ST11 isolates grouped with strains from the globally disseminated clonal group (CG) 258 (also including ST258, ST340, ST437, and ST512), comprising most CR-*Kp* isolated worldwide and highly represented in the NCBI database. The other isolates covered less represented groups, outstanding ST505, corresponding to only five genomes from our set.

**Figure 1.**
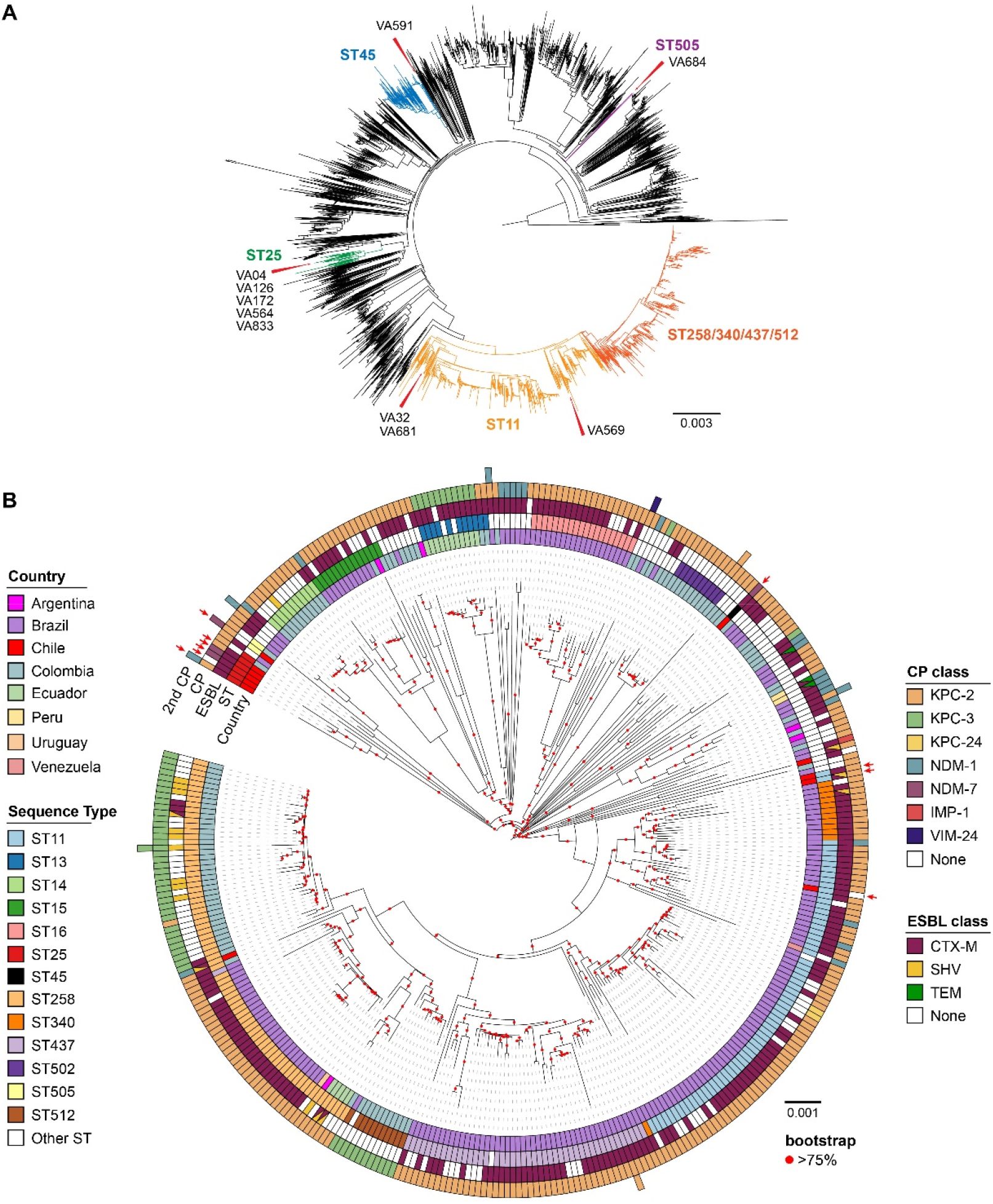
Phylogenomic relationships between Chilean CR-*Kp* isolates and *K. pneumoniae* isolated worldwide. (A) Phylogenetic tree including the Chilean isolates (red arrows) and 3443 Kp1 genomes retrieved from the NCBI database. Branches corresponding to relevant sequence types (STs) are colored. (B) Phylogenetic tree with 353 *K. pneumoniae* strains encoding carbapenemases isolated in South America. The node color indicates bootstrap values. The tracks correspond to the country of origin, sequence type, and the kind of one or possibly two carbapenemase genes.

**Table 2.** Genome assembly stats.

| Strain | ST | Total length (bp) | %GC | Cov. depth | Compl. (%) | Contam. (%) | Chromosome length (bp) | N° plasmids | Plasmids length (kbp) | Accession |
| --- | --- | --- | --- | --- | --- | --- | --- | --- | --- | --- |
| VA04 | ST25 | 5,762,453 | 56.98 | 176X | 99.90 | 0.90 | 5,456,744 | 4 | 6*; 46; 58; 196 | CP093501-CP093505 |
| VA32 | ST11 | 5,731,565 | 57.17 | 202X | 100.00 | 0.08 | 5,286,512 | 5 | 7; 58; 74; 105; 200 | CP093495-CP093500 |
| VA126 | ST25 | 5,757,426 | 56.98 | 181X | 92.28 | 0.90 | 5,121,870 | 4 | 12; 42; 68; 175 | CP093489-CP093494 |
| VA172 | ST25 | 5,403,938 | 57.44 | 198X | 100.00 | 0.39 | 5,246,520 | 2 | 67; 90 | CP093486-CP093488 |
| VA564 | ST25 | 5,718,217 | 57.00 | 182X | 100.00 | 1.21 | 5,406,546 | 4 | 33; 37; 63; 179 | CP093481-CP093485 |
| VA569 | ST11 | 5,713,876 | 57.13 | 164X | 100.00 | 0.59 | 5,584,676 | 4 | 6; 7; 35; 78 | CP093476-CP093480 |
| VA591 | ST45 | 5,653,497 | 57.18 | 180X | 100.00 | 0.08 | 5,430,590 | 3 | 9; 46; 160 | CP093471-CP093475 |
| VA681 | ST11 | 5,554,568 | 57.14 | 204X | 99.63 | 0.39 | 5,250,844 | 5 | 5; 14; 36; 58; 191 | CP093465-CP093470 |
| VA684 | ST505 | 5,818,826 | 56.74 | 195X | 100.00 | 0.08 | 5,176,823 | 7 | 2; 6; 8; 37; 49; 146; 388 | CP093458-CP093464 |
| VA833 | ST25 | 5,997,469 | 56.80 | 220X | 99.74 | 1.25 | 5,458,013 | 6 | 7; 45*; 56*; 92; 165; 176 | CP093451-CP093457 |

Next, we conducted a core-genome MLST analysis to examine the clonality among the isolates and 55 strains showing the lowest Mash distance to them, clustering into four clonal groups, as defined by Bialek-Diavenet et al. (2014)[30] (Supplementary Figure 1). Most isolates appeared mixed with others from diverse geographic origins and thus would correspond to disseminated clones, while VA591 formed a separate branch inside CG45, apparently more constrained to Chile.

To place the Chilean isolates in a regional CR-*Kp* landscape, we restricted the Mashtree analysis to 351 isolates encoding carbapenemases from South America, mainly from Brazil (215) and Colombia (108), plus other countries considerably less represented in the database. Also, for comparison, we included two draft genomes of Chilean *Kp* isolates lacking carbapenemases published previously[31]. Thus, we observed more than 55 STs that tended to cluster following the geographical origin, predominating ST258 from Colombia and ST11 and ST437 from Brazil (Figure 1B). Further, the Chilean ST25 isolates clustered together, close to the only other ST25 isolate (from Colombia). Similarly, the ST505 isolate shared a deep branch with one from Colombia, the only two ST505 isolates encoding carbapenemases found in the NCBI assembly database. On the other hand, VA591 formed a separate branch and was the only South American carbapenemase-producing ST45 isolate. Furthermore, two Chilean ST11 formed a clade with ST340 isolates from Brazil and the third with ST11 from the same country. Also, the previously described Chilean genomes grouped apart from the isolates described in this study.

Regarding beta-lactamase gene content, most South American isolates encoded one carbapenemase, predominating KPC-2, followed by KPC-3 and NDM-1 (Figure 1B). Additionally, nine isolates had two carbapenemase genes (mainly KPC-2 plus NDM-1), including VA684 and VA833. Moreover, 218 isolates (62%) carried one or more acquired ESBLs, mostly CTX-M-15, CTX-M-2, and CTM-X-14, with the sporadic presence of SHV-12, TEM-26, and TEM-15.

### Genome-wide resistance genes profile

Using the complete genome information, we identified the antibiotic-resistance genes of the Chilean isolates and compared the prevalence of their beta-lactamases (and combinations) with those found in the 3443 genome set. Following the high experimental beta-lactam resistance, we identified up to eight beta-lactamase genes per isolate (Figure 2A, Supplementary Table 4 and 5), three carrying KPC-2, the most common carbapenemase among our set, and one KPC-2+NDM-1, a relatively rare combination (Supplementary Figure 2A). Moreover, three isolates had NDM-7, a significantly less frequent variant of NDM-1 with increased activity, while VA684 was the only genome from the set carrying NDM-7+KPC-2.

**Figure 2.**
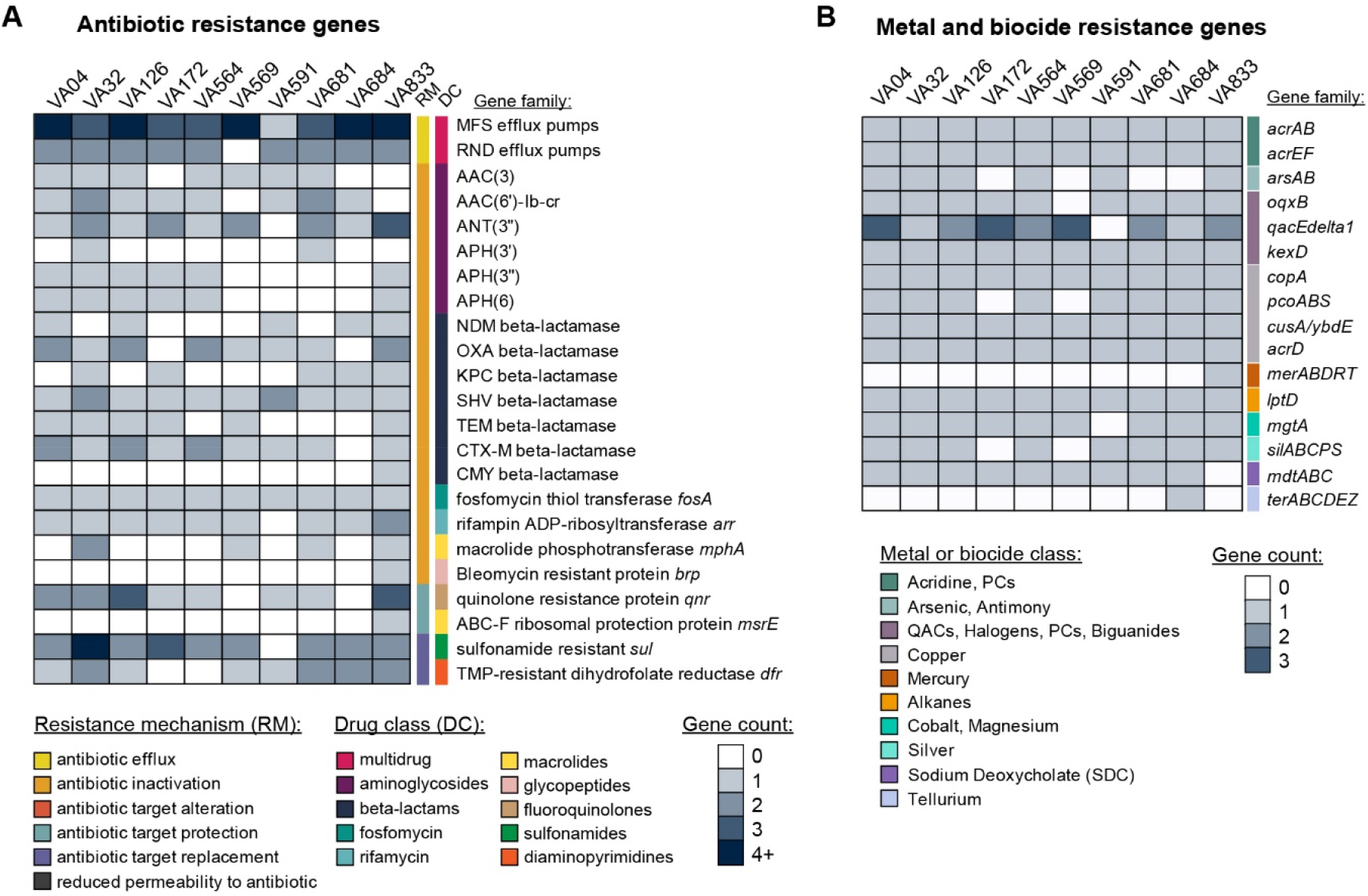
Genome-wide resistance gene profile of the CR-*Kp* isolates. Bioinformatic prediction of genes conferring resistance to antibiotics (A), or to metals, disinfectants, and other biocides (B). The heatmap color scale indicates the number of genes found per genome in each category.

Additionally, we identified CTX-M-15 and the less frequent CTX-M-2 and CTX-M3 ESBLs (Supplementary Figure 2B) among our isolates, three encoding the CTX-M-15+CTX-M-2 and one the CTX-M-15+SHV-31 rare combinations. Other acquired beta-lactamases included the rare combinations OXA-1+OXA-10 (VA564) and OXA-1+OXA-10+TEM-1+CMY-2 (VA833). Of note, the CMY-2 class C beta-lactamase is relatively rare in *K. pneumoniae* and was absent in the rest of the 353 Latin American *K. pneumoniae* analyzed. Additionally, six isolates had mutations in the OmpK35 and OmpK36 porins, shown to confer carbapenem resistance in other *K. pneumoniae* strains, which could explain the carbapenem resistance in VA564 and VA569 lacking carbapenemase genes.

Besides, several aminoglycoside resistance genes were found, mainly ANT(3’’)-IIa, AAC(3)-IIe, AAC(6’)-lb-cr (also reported to confer fluoroquinolone resistance[32]), APH(3’’)-Ib, and APH(6)-Id (Figure 2A, Supplementary Table 4 and 5). Accordingly, most isolates were resistant to gentamicin (Table 1). On the other hand, VA833 was the only isolate carrying the *armA* gene encoding a 16S rRNA (guanine(1405)-N(7))-methyltransferase, which could account for its unusually high amikacin resistance (Table 1), as reported previously in *K. pneumoniae* [33].

Regarding fluoroquinolone resistance, most isolates encoded the *adeF* and *oqxA* efflux pumps, up to four quinolone resistance proteins (mainly QnrB1 and QnrB19), and carried one or two point mutations in *gyrA* (mainly 83Y, 83I, 87G, 87N) and the *parC*-80I mutation (Figure 2A, Supplementary Table 4). These findings correlated with ciprofloxacin resistance in most isolates, while the absence of acquired fluoroquinolone resistance genes and mutations in target host genes could explain VA684 sensitivity.

Regarding colistin resistance, none of the isolates were positive for *mcr-1* or *mcr-2* genes. Conversely, all the isolates were positive for *eptB* and *arnT* genes encoding phosphoethanolamine transferases, the only identified genes linked to peptide antibiotics resistance. Furthermore, only *tetA* and *tetD* genes linked to tetracycline resistance were found in five isolates, including VA04 and VA833. Thus, no clear genetic bases for the high tigecycline resistance shown by these two isolates, nor for the VA564 and VA569 colistin resistance, could be established.

Among other resistance determinants found in the Chilean strains, mainly in VA833, are the macrolide 2’-phosphotransferase MPH(2’)-I and a BRP(MBL) bleomycin resistance protein. On the other hand, most isolates encoded several genes involved in resistance to metals and biocides, including copper, arsenic, silver, mercury, tellurium, acridine, halogens, phenolic and quaternary ammonium compounds, and bile salts (Figure 2B).

### Virulence factors present in Chilean CR-Kp

Next, we identified and categorized the virulence factors present in the Chilean CR-Kp (Figure 3). The five ST25 isolates displayed a K2 capsule associated with increased virulence[2], while the ST11 isolates had KL15 and KL39. The other isolates had KL24 (ST45) and KL64 (ST505). In terms of adhesion factors, all had the *fim, mrk, ecp*, and *pil* operons, encoding a type 1 (T1P) fimbriae, a type 3 (MR/K) pili, a homolog of the *E. coli* common pilus (ECP), and a type 4 hemorrhagic *E. coli* pilus (HCP), respectively, all of them reported to be involved in cell adherence to tissues [34]. Further, VA684 has a homolog of the type 4 *stb* fimbrial operon linked to long-term intestinal carriage in *Salmonella* Typhimurium [35].

**Figure 3.**
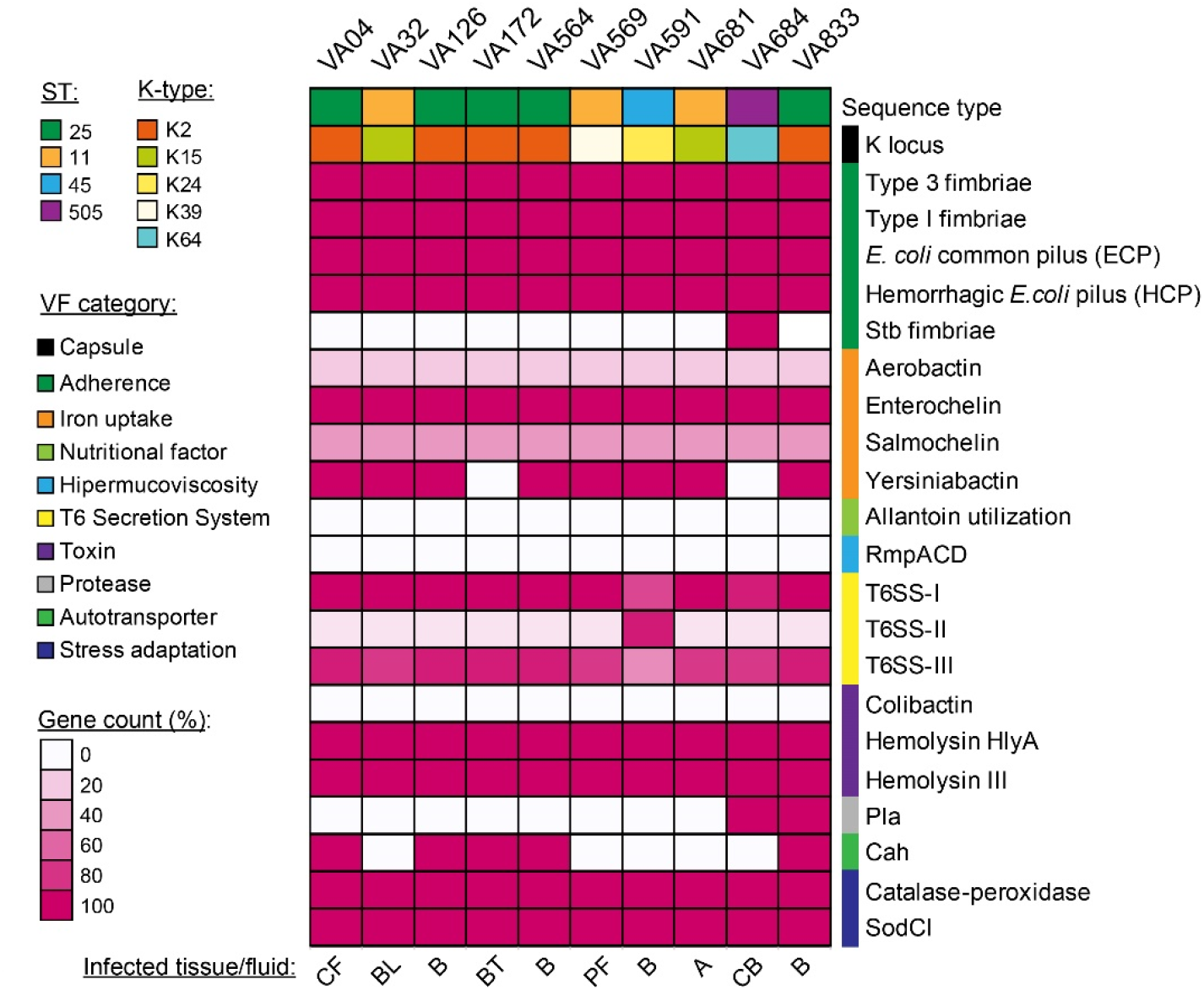
Pathogenic potential encoded in the Chilean CR-*Kp*. Bioinformatic prediction and categorization of genes linked to virulence. The heatmap colors represent the percentage number of genes involved in the respective virulence trait found in each isolate over the number of known genes required to express that trait. CF: cerebrospinal fluid, BL: bronchoalveolar lavage, B: blood, BT: bone tissue, PF: peritoneal fluid, A: abscess, CB: catheter blood.

Regarding siderophore systems, all or most isolates had the *ent* and *ybt* genes for enterobactin and yersiniabactin production, while none of them would produce aerobactin and salmochelin, which determinants are frequently encoded in the large *Klebsiella* virulence plasmids (pKpVPs)[2]. Moreover, all the strains lacked the hypermucoviscosity-related genes *rmpACD*, which are also typically found in these plasmids. Regarding cell-to-cell killing capacity, all the isolates had two type-VI secretion systems (T6SS-I and T6SS-III), while VA684 also had a third system (T6SS-II). Finally, in terms of toxin production, all the isolates encoded the hemolysin-like proteins HlyA and hemolysin III, while none showed the determinants for colibactin production, which was strongly associated with hypervirulent strains.

Other factors present in some strains were the Pla protease from *Yersinia pestis*, which prevents host cell apoptosis and inflammation by degrading Fas ligand (FasL) [36], and the Cah autotransporter associated with enterohemorrhagic *E. coli* infections [37]. Thus, we identified an ample array of virulence factors in the Chilean *K. pneumoniae* isolates that would contribute during infection, which, however, lack the hallmark factors associated with the hypervirulent pathotype.

### Plasmids encoding carbapenemases and other resistance determinants

We examined 44 contigs identified as plasmids among the ten isolates (39 circular replicons and five fragments) (Supplementary Table 6). After annotation, 27 plasmids were predicted as conjugative (oriT + T4SS), 14 mobilizable (oriT only), and three lacked conjugation determinants. In addition, over ten compatibility groups were observed, predominating IncR, IncFIB(K), and IncFII(K). Upon sequence-based clusterization, nine groups and 15 singletons were observed, with an overall limited number of shared plasmids even among closely related isolates, except for VA04 and VA126 (ST25) harboring four from the same group. Moreover, intra-cluster sequence comparisons revealed substantive smaller-scale variability (Supplementary Figure 3).

Ten conjugative plasmids encoding carbapenemases were identified, four NDM-7, five KPC-2, and one NDM-1 (Figure 4). The NDM-1 plasmid pVA833-165 (d~165-kbp, IncCc) showed 99% identity and 89% coverage with an uncharacterized plasmid from *Salmonella enterica* (CP009409) and ~88% coverage with an IncA/C plasmid carrying a blaVIM gene from *K. pneumoniae* ST383 (KR559888.1). However, pVA833-165 differed in carrying the blaNDM-1 gene inside an IS3000-formed composite transposon, along with blaCMY-2, *floR, tet(A), APH(6)-Id, sul2*, and the mercury resistance genes *merRTPABDE* (Supplementary Figure 4A), pointing out the high-risk potential of this plasmid.

**Figure 4.**
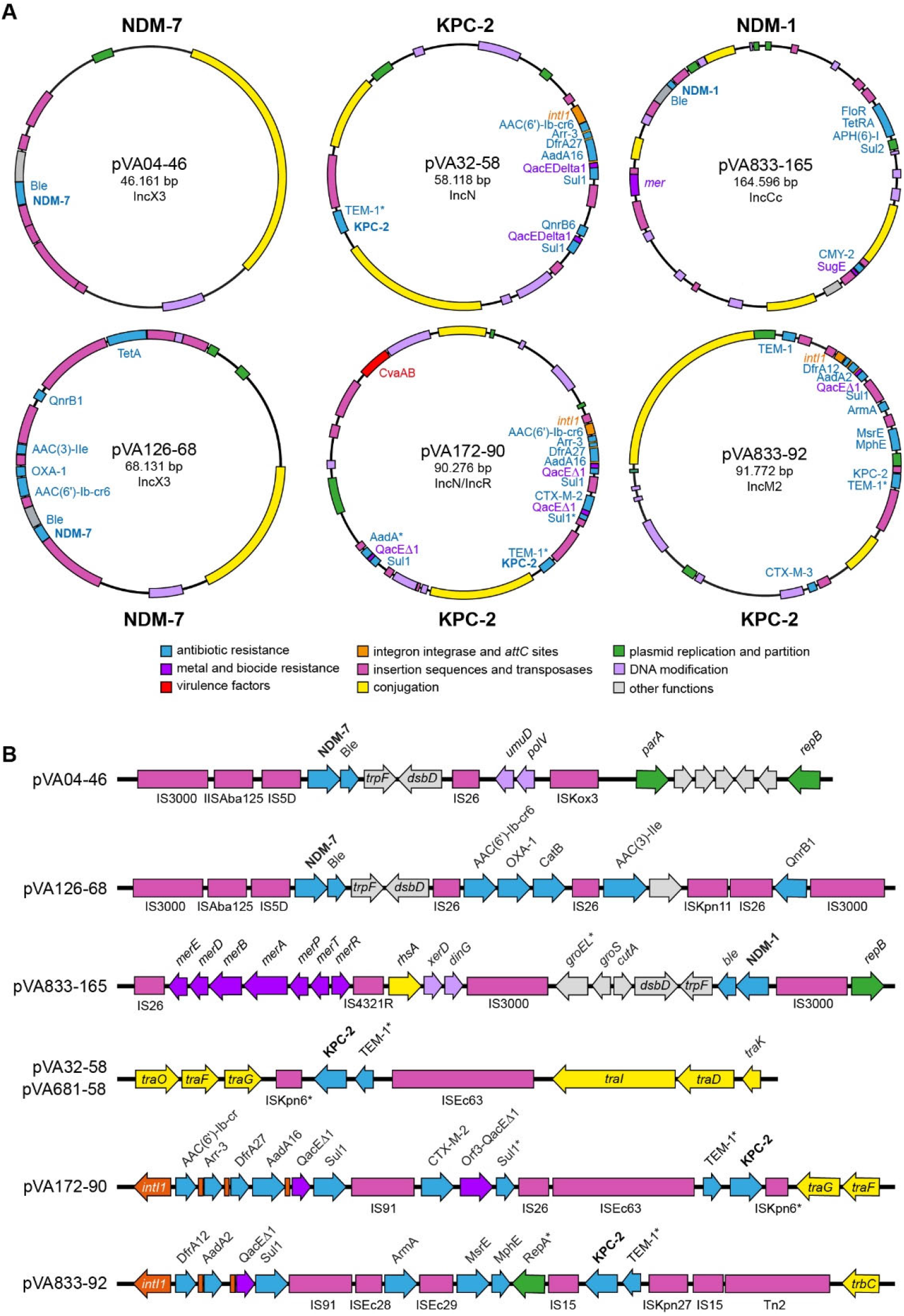
Plasmids carrying carbapenemase genes and other resistance determinants. (A) Schematic representation of the main NDM-1, NDM-7, and KPC-2 plasmids and their encoded functions. (B) genomic context of the genes encoding carbapenemases and other resistance determinants.

The NDM-7 plasmids pVA04-46, pVA591-46, pVA684-49, and pVA126-68 (IncX3), showed ~99% identity and coverage with 46-kbp plasmids from the NCBI database, including pJN05NDM7 (NZ_MH523639.1) from uropathogenic *E. coli*. In these plasmids, the blaNDM-7 gene is located in the genetic context described originally[38] (Supplementary Figure 4B). Remarkably, pVA126-68 differed from other NDM-7 plasmids, in bearing an extra 21-kbp region comprising a CALIN element (clusters of attC sites lacking integron-integrases) associated with AAC(6’)-Ib-cr, OXA-1, CatB, and AAC(3)-IIe, plus other resistance determinants (Figure 4).

Additionally, the KPC-2 plasmids pVA172-90, pVA32-58, pVA681-58, and pVA684-37 (IncN) were ~100% identical, except for the former showing additional segments. The *bla*_KPC-2_ gene was found in the NTE_KPC_-IIe genetic environment (Supplementary Figure 5A), first identified in IncN plasmids from a recent Chilean outbreak of carbapenem-resistant bacteria (e.g., pKpn-3; MT949189.1)[39] and in p33Kva16-KPC and pEC881_KPC from Colombian *E. coli* and *K. variicola* isolates[40], showing high identity with pVA681-58 and related plasmids (Supplementary Figure 5B). These KPC plasmids included a class-I integron carrying AAC(6’)-Ib-cr6, *arr-3, dfrA27*, and *aadA16*, and a region encoding QnrB6, QacEDelta1, and Sul1. In pVA172-90, this last region was replaced by one including *bla*_CTX-M-2_ and other resistance genes. Also, it has a ~25-kbp extra region including *sul1, qacEdelta1*, and the CvaAB hemolysin exporter, among other features. Furthermore, pVA172-90 showed ≤57% coverage matches in the NCBI database, thus would be a novel KPC-2 plasmid variant with additional resistance determinants.

A further plasmid bearing KPC-2, differing considerably from the others, was pVA833-92 (IncM2). This plasmid showed 97% coverage and 99% identity with pCTX-M-3 from *Citrobacter freundii* (Supplementary Figure 6A) encoding the CTX-M-3 ESBL, *bla*_TEm-1_, *aacC2*, and *armA*, as well as an integron carrying *aadA2, dfrA12*, and *sul1*. Strikingly, pCTX-M-3 lacked the KPC-2 gene found in pVA833-92. The inspection of the *bla*_KPC-2_ genetic environment revealed that it is immediately downstream of *bla*_TEM-1_ (truncated) and ISKpn27 (formerly ISKpn8) (Supplementary Figure 6B). Two IS26 elements that flank these three genes could have mediated the insertion of this segment into a pCTX-M-3-like plasmid. This context differed from all the known NTE_KPC_ environments (Supplementary Figure 6C) and would correspond to a novel genetic platform for mobilizing *bla*_KPC-2_, tentatively NTE_KPC_-IIf.

Besides those encoding carbapenemases, we found other relevant plasmids (Supplementary Figure 7). pVA04-196 (IncFIB(K)/IncFII(K)) outstood for carrying three metal resistance operons (*sil*, *pco*, and *ars*), TEM-1, OXA-1, and a CTX-M-15. pVA32-200 (Col440I/IncFIB(K)/IncFII(K)) also had the *sil, pco*, and *ars* operons, along with a wide array of antibiotic resistance genes and determinants for Klebicin B production. Finally, we found plasmids potentially contributing to bacterial fitness during the infective process. For example, pVA684-388 (IncR, ~388-kbp) encoded determinants for K+ uptake, trehalose synthesis, putrescine utilization, molecular chaperones, heat shock proteins metabolism, and glycerol utilization.

### Genomic islands encoding virulence and fitness factors

Besides plasmids, genomic islands (GIs) and prophages are vehicles for disseminating clinically relevant traits in pathogens. In this regard, the chromosome alignment among the Chilean isolates showed different variable regions (Figure 5A), including putative MGEs integrated into transfer RNA genes (tDNAs), which act as hotspots for GI and prophage integration in *Klebsiella* and other bacteria [41].

**Figure 5.**
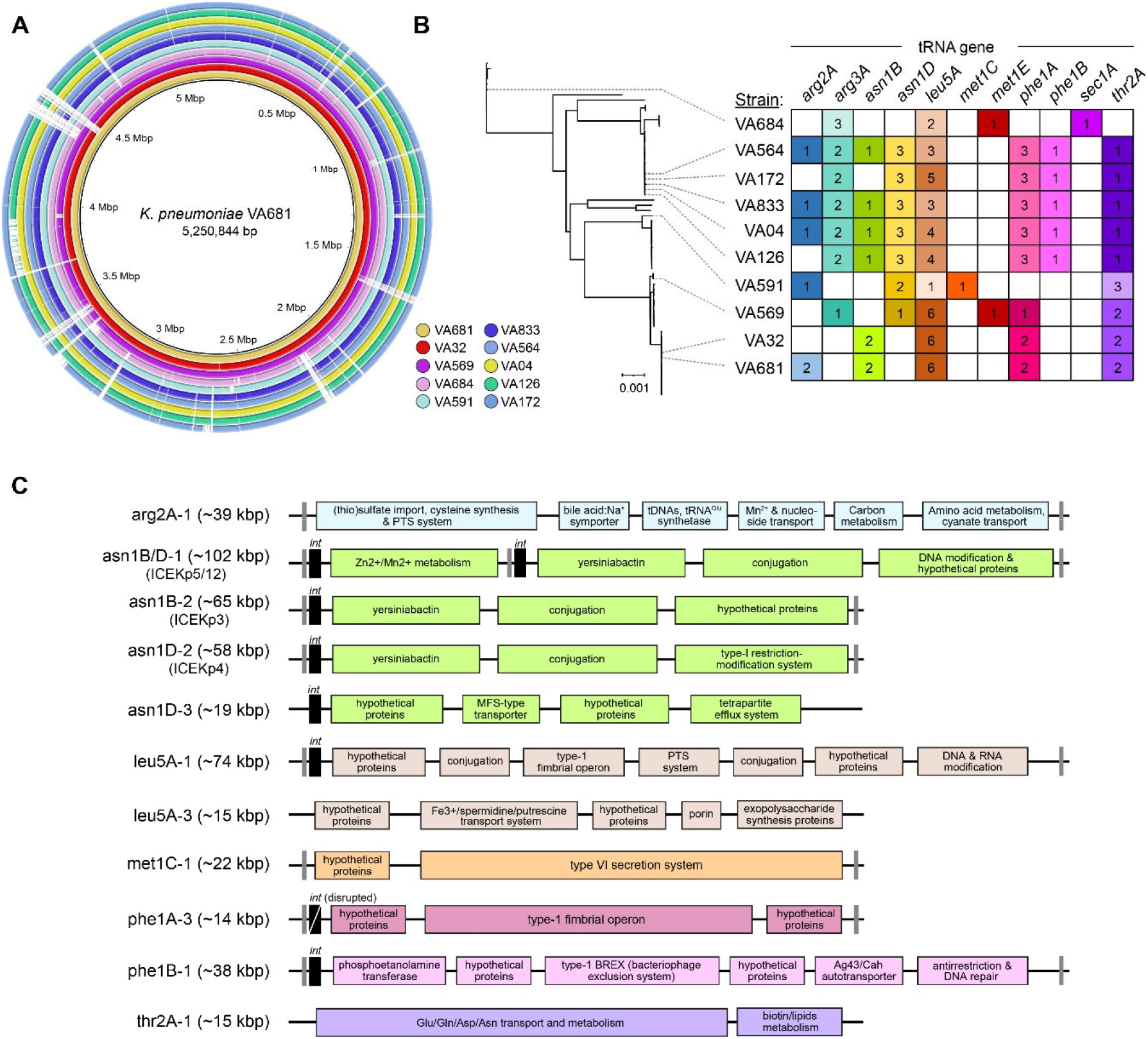
Chromosomal mobile genetic elements and encoded functions. (A) Chromosome sequence alignment among the Chilean isolates. (B) Mobile genetic elements integrated into tRNA genes. The numbers indicate different MGEs. The distance tree was built based on cgMLST analysis. (C) Genomic islands encoding virulence and fitness factors. The gray bars indicate direct repeats; *int*: integrase.

Following a pipeline for GI and prophage prediction among the isolates, we identified 61 MGEs (24 non-redundant) integrated into 11 tDNAs, with sizes ranging from ~7.5 to ~103 kbp. Furthermore, a variable chromosomal MGE content was observed even among closely related strains (i.e., ST25 and ST11), especially in *leu5A, asn1B*, and *arg2A* loci, suggesting that active traffic of GIs and prophages would contribute to chromosomal variability in Chilean CR-*Kp* isolates.

After annotation, eight of the twenty-four non-redundant MGEs corresponded to intact, incomplete, or questionable prophages, while the rest were classified as GIs, most encoding a P4-like integrase and flanking direct repeats (Supplementary Table 7). Twelve of these GIs encoded factors potentially linked to pathogenesis or bacterial fitness, including three members of the ICEKp family (ICEKp3/5/12) integrated into asparagine tDNAs (Figure 5C). These elements are widely disseminated *in K. pneumoniae* and carry genes for yersiniabactin siderophore production and additional variable cargo modules[42]. Additionally, we identified novel GIs encoding relevant functions, including carbon, amino acids, and bile acids metabolism and transport, the Cah autotransporter, fimbrial operons, a Fe3+/spermidine/putrescine transport system, and a type-VI secretion system, among others. Thus, different GIs would mediate virulence and fitness factors dissemination among Chilean CR-*Kp*.

## DISCUSSION

Genome analysis is a powerful approach to support pathogen surveillance, although it is still restricted in lower and middle-income countries due to economic limitations and less expeditious access to equipment and supplies [43]. Regrettably, this causes many geographical regions, including most of South America, to be underrepresented in genome databases and, thus, elusive in the global epidemiological landscape. This study presented the first ten complete genome sequences of CR-*Kp* causing infections in Chile, combining this information with phenotypic tests to unveil their resistance mechanisms and pathogenic potential, providing relevant baseline knowledge for further local and global studies on *K. pneumoniae* epidemiology.

Although most CR-*Kp* infections worldwide were attributed to CG258 and a few other lineages, the burden of local problem clones remains substantial [2]. CG258 KPC-producing *Kp* has been widely reported in South America, although many less frequent STs were also found [4,44–46]. In Chile, reported CR-*Kp* mainly belong to ST11, ST258, ST29, ST1588, and ST1161 (potentially native to Chile) [47,48]. Conversely, ST25, linked to a relatively low number of CR-*Kp* infections worldwide predominated in this study, as previously reported in Argentina [49]. Also, ST25 CR-*Kp* causing invasive infections were isolated from an outbreak in Ecuador [50]. As in this study, these ST25 isolates showed a K2 capsular serotype linked to increased virulence [2], although no genomic data is available for comparison. Despite two draft genomes of isolates from Argentina proposed to be ST25 were published [51], our analyses indicated that they actually are ST629 and ST551-1LV.

Thus, ST25 K2 CR-*Kp* seems to be an emerging clone in South America, where carbapenem resistance and increased virulence converge, and KPC-2 and NDM-1 are by far the most common carbapenemases. However, four isolates described here encoded the NDM-7 carbapenemase, corresponding to the first report of NDM-7 ST25 CR-*Kp* in South America, further stressing the rapid evolution and potential risk of this lineage.

Additionally, we identified two isolates from the less frequent ST45 and ST505. ST45 is an emerging carbapenemase-producing clone highly prevalent in the EuSCAPE surveillance study [14,52]. Also, ST45 KPC-2-producing *Kp* were reported in Venezuela [53], while infections of ST45 *Kp* lacking carbapenemases were reported in Brazil and Mexico [54,55]. Thus, to our knowledge, VA591 corresponds to the first report of ST45 *Kp* producing NDM-7 in South America, which also showed resistance to colistin and diverged from other ST45 isolated worldwide. On the other hand, very few studies described ST505 *Kp* infections, including one from Mexico[54], although these isolates were carbapenem-sensitive. We found no previous reports of ST505 CR-*Kp* and VA684 was the only ST505 isolate from our genome database producing carbapenemases. Thus, this would be the first report of an ST505 *Kp* producing both NDM-7 and KPC-2 worldwide.

The high levels of resistance found in the isolates correlated with an ample array of acquired resistance genes, some found with relatively low frequency in other *K. pneumoniae* isolates, including *bla*NDM-7, alone or combined with *bla*KPC-2 (first report in Chile), *bla*SHV-31 (first report in Chile), and the CMY-2 ampC beta-lactamase (first report in *K. pneumoniae* in Chile). Moreover, most isolates also carried genes conferring resistance to metals and disinfectants, in agreement with previous reports showing their co-selection with antibiotic resistance genes, widening the toolbox to resist antimicrobial chemotherapy and persist in clinical, urban, and polluted environments [56,57].

The characterization of the mobile elements mediating the dissemination of clinically relevant traits among the Chilean isolates led to the identification of numerous plasmids and GIs, anticipating a remarkable diversity of these elements in the *K. pneumoniae* population. The sequencing of additional genomes, especially from underrepresented geographical areas, will contribute to unveiling this yet-to-be-known MGE diversity. Most of these MGEs included resistance and virulence genes, but also genes potentially involved in cell fitness, whose contribution to host infection deserves further studies.

Plasmids carrying carbapenemase genes, specially *bla*KPC, are the main drivers of carbapenem resistance in *Enterobacteriaceae* globally [58]. Although *bla*KPC genes were initially associated with a ~10 kb Tn4401 derivative, novel non-Tn4401 elements (NTEKPC) for their dissemination have been described [58]. Both platforms were described in a previous study in Chile [47], with the predominance of the so-called Argentinian variant 1a [59] (NTEKPC-IIa) in non-conjugative plasmids from ST11, ST29, and ST1161 *Kp*. Conversely, in this study, pVA32-58 and related conjugative plasmids (IncN) from ST25, ST11, and ST505 isolates, had the *bla*KPC-2 gene in the NTEKPC-IIe genetic environment, which recently emerged in Chile in an ST11 *K. pneumoniae* (and other *Enterobacteriaceae)*, and in *K. variicola* in Colombia [39,40]. Moreover, pVA172-90 was a new variant with additional resistance genes, including *bla*CTX-M-2. On the other hand, to our knowledge, pVA833-92 corresponds to the first report where *bla*KPC-2 jumped to a pCTX-M-3-like IncM2 plasmid, located in a novel class II KPC genetic environment flanked by IS26 elements named NTEKPC-IIg.

Although KPC is the most prevalent carbapenemase in *K. pneumoniae* worldwide, half of the isolates studied here encoded an NDM carbapenemase, two showing the co-existence of both enzymes. In Chile, *K. pneumoniae* producing NDM-1 was first reported in 2018 in an ST1588 isolate [48]. The novel *bla*_NDM-1_ plasmid pVA833-165 (IncC) from an ST25 isolate had a known IS3000 environment but differed from known plasmids in carrying *bla*_CMY-2_ and several other antibiotic and metal resistance genes. Meanwhile, the *bla*NDM-7 plasmids (IncX3) had a Tn125 environment, where pVA126-68 differed from previously known plasmids in having an additional 21-kbp region encoding several additional resistance genes.

On the other hand, the genome-based virulence profiling revealed several known factors linked to cell adhesion, iron uptake, and cell-to-cell killing. Also, other factors not described before in *K. pneumoniae*, such as the Cah autotransporter from enterohaemorrhagic *E. coli* (EHEC) involved in autoaggregation and biofilm formation, and the *Y. pestis* Pla protease involved in signaling to manipulate host cell death and inflammation. Further studies are required to investigate the role of these factors in *K. pneumoniae* pathogenesis.

Of special concern are hypervirulent *K. pneumoniae* strains causing pyogenic liver abscesses and other severe community-acquired infections, mainly in Southeast Asia. Although initially some studies defined hypervirulence by a positive string test indicating hypermucoviscosity, it is now accepted that is more appropriate to combine aerobactin positivity and *rmpADC* gene presence as markers of hv*Kp*, which are strongly associated with ST23 and K1/K2 capsule [2]. None of the isolates described here have these hv*Kp* hallmark features. A previous work reported the isolation of an ST1161 hypermucoviscous CR hv*Kp* strain in Chile[31], however, it lacked *rmp* and aerobactin production genes, had a K19 capsular serotype, and lacked the colibactin synthesis genes also commonly found in hv*Kp*. A similar situation occurred in a ST29 CR hvKp report from Brazil [60]. In both cases the virulence was assayed using a *Galleria mellonella* infection model, which has limited utility to distinguish classical virulent *Kp* from hv*Kp*. Thus, more studies are required to confirm the presence of hv*Kp* in Chile and South America.

## ACKNOWLEDGEMENTS

This study was supported by grants FONDECYT 11181135 and 1221193 (A. Marcoleta), FONDECYT 1211852 (F. Chávez) and the Millenium Institute Center for Genome Regulation (M. Allende). Also, it was supported by Instituto de Salud Pública de Chile.

