## Supplementary Material for "Population genomics, resistance, pathogenic potential, and mobile genetic elements of carbapenem-resistant *Klebsiella pneumoniae* causing infections in Chile"

**Supplementary Table 1.** *K. pneumoniae* isolates studied in this work.

| Isolate | Sex | Age | Collection date | Source | Region of origin |
|---|---|---|---|---|---|
| VA4 | M | 36 | 05-12-2018 | Cerebrospinal fluid | Metropolitan |
| VA32 | F | 81 | 04-01-2019 | Bronchoalveolar lavage | Metropolitan |
| VA126 | M | 33 | 16-01-2019 | Blood | Metropolitan |
| VA172 | M | 71 | 25-01-2019 | Bone tissue | VII Del Maule |
| VA564 | M | 79 | 02-04-2019 | Blood | X De los Lagos |
| VA569 | M | 53 | 06-04-2019 | Peritoneal fluid | Metropolitan |
| VA591 | M | 29 | 11-04-2019 | Blood | Metropolitan |
| VA681 | F | 32 | 24-04-2019 | Abscess | Metropolitan |
| VA684 | F | 15 | 25-04-2019 | Catheter blood | Metropolitan |
| VA833 | F | 54 | 23-05-2019 | Blood | Metropolitan |

**Supplementary Table 2.** Phenotypic tests for carbapenemase production and beta-lactamase gene detection in the *K. pneumoniae* isolates studied in this work.

| Isolate | Blue Carba | Boronic Acid | Triton Hodge | carbapenemase gene | ESBL |
|---|---|---|---|---|---|
| VA4 | + | - | + | NDM+ | + |
| VA32 | + | + | + | KPC+ | + |
| VA126 | + | - | + | NDM+ | + |
| VA172 | + | + | + | KPC+ | + |
| VA564 | - | - | - | - | + |
| VA569 | - | - | - | - | + |
| VA591 | + | - | + | NDM+ | + |
| VA681 | + | + | + | KPC+ | + |
| VA684 | + | + | + | KPC+/NDM+ | + |
| VA833 | + | + | + | KPC+/NDM+ | + |

**Supplementary Table 3.** List and accessions of the 3443 *K. pneumoniae* genomes included in the analyses presented in this work. (Provided as a separate spreadsheet).

**Supplementary Table 4.** Harmonized list of antibiotic resistance genes identified in the Chilean *K. pneumoniae* isolates described in this study. (Provided as a separate spreadsheet).

**Supplementary Table 5.** Summary of the main antibiotic resistance determinants identified per isolate and drug class. (Provided as a separate spreadsheet).

**Supplementary Table 6.** Main features of the plasmids identified in the *K. pneumoniae* isolates described in this study (Provided as a separate spreadsheet).

**Supplementary Table 7.** tDNA-associated mobile genetic elements found in the chromosome of the *K. pneumoniae* isolates described in this work (Provided as a separate spreadsheet).

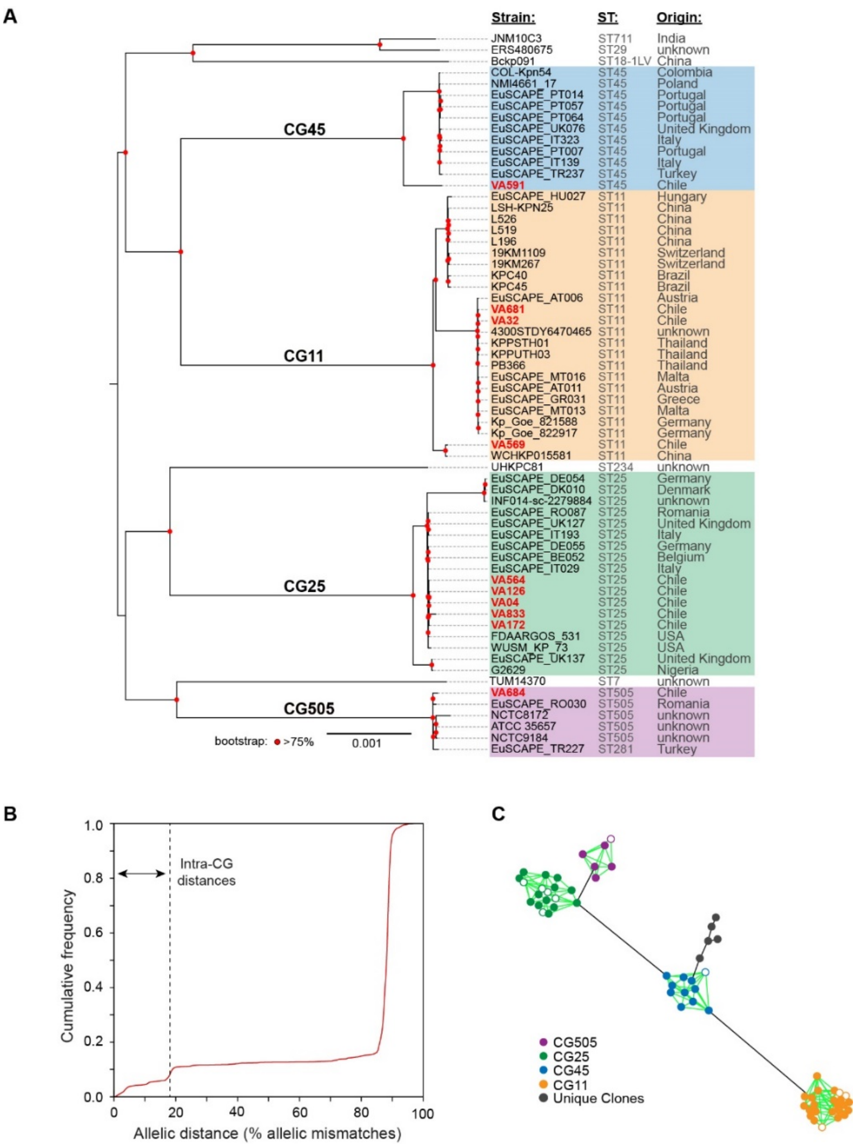


**Supplementary Figure 1. The ten Chilean CR-Kp isolates described in this work fall into five well-defined clonal groups.** (A) cgMLST-based distance tree, including the isolates described in this work and 55 closely related strains from our 3443 *K. pneumoniae* genome set, based on Mash distances. The colors indicate different STs, following Kleborate tool prediction. The color nodes represent the result of 1000 bootstrap iterations. (B) Frequency distribution plot of the allelic distances among the 3443 *K. pneumoniae* genomes used to estimate the 18.4% distance limit for clonal groups, based on a strategy described by Bialek-Davenet et al. (2014) [18]. (C) Minimum expansion tree showing the clusters formed by genomes from the tree shown in (A) having a distance <18.4% and thus forming part of the same clonal group (shown in the same color) Circles with white fill represent the Chilean isolates described in this study. We observed a clear separation between intra- and inter-group allelic distances, supporting a <18% allelic mismatches boundary for clonal groups. To calculate allelic distances, the set of 3453 genomes was used as input for the chewBBACA v2.7.0 pipeline. Using the default parameters and a 95% gene presence, we created a 2,750 loci cgMLST scheme using 639 high-quality Kp1 assemblies (N50>5Mbp, no Kleborate QC warnings), which was then applied to the other genomes. The typing matrix obtained was then used as input for the PHILOViZ online visualization tool to generate the minimum expansion tree. Additionally, the scheme was used to calculate the allelic distances between all genome pairs and construct a histogram to establish coherent limits for intra-CG distances.

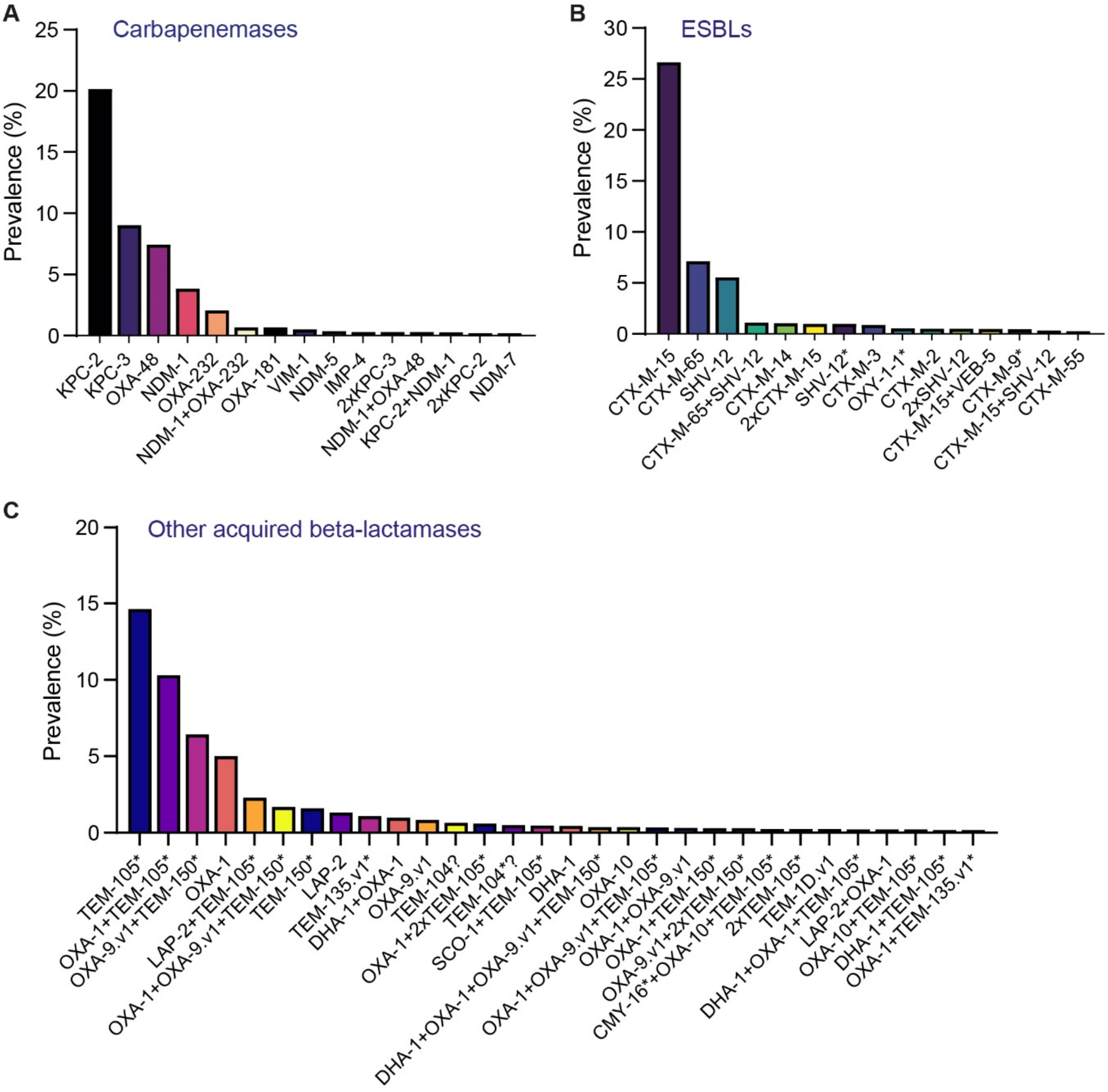


**Supplementary Figure 2.** Prevalence of genes encoding carbapenemases (A), ESBLs (B), and other acquired beta-lactamases (C) found in our 3443 *K. pneumoniae* genome set according to Kleborate prediction. (*): no exact amino acid match was found; the closest nucleotide match is reported instead. (?) truncated protein compared with the reference database. The prevalence was determined as the number of genomes showing each beta-lactamase combination divided by the total number of genomes.

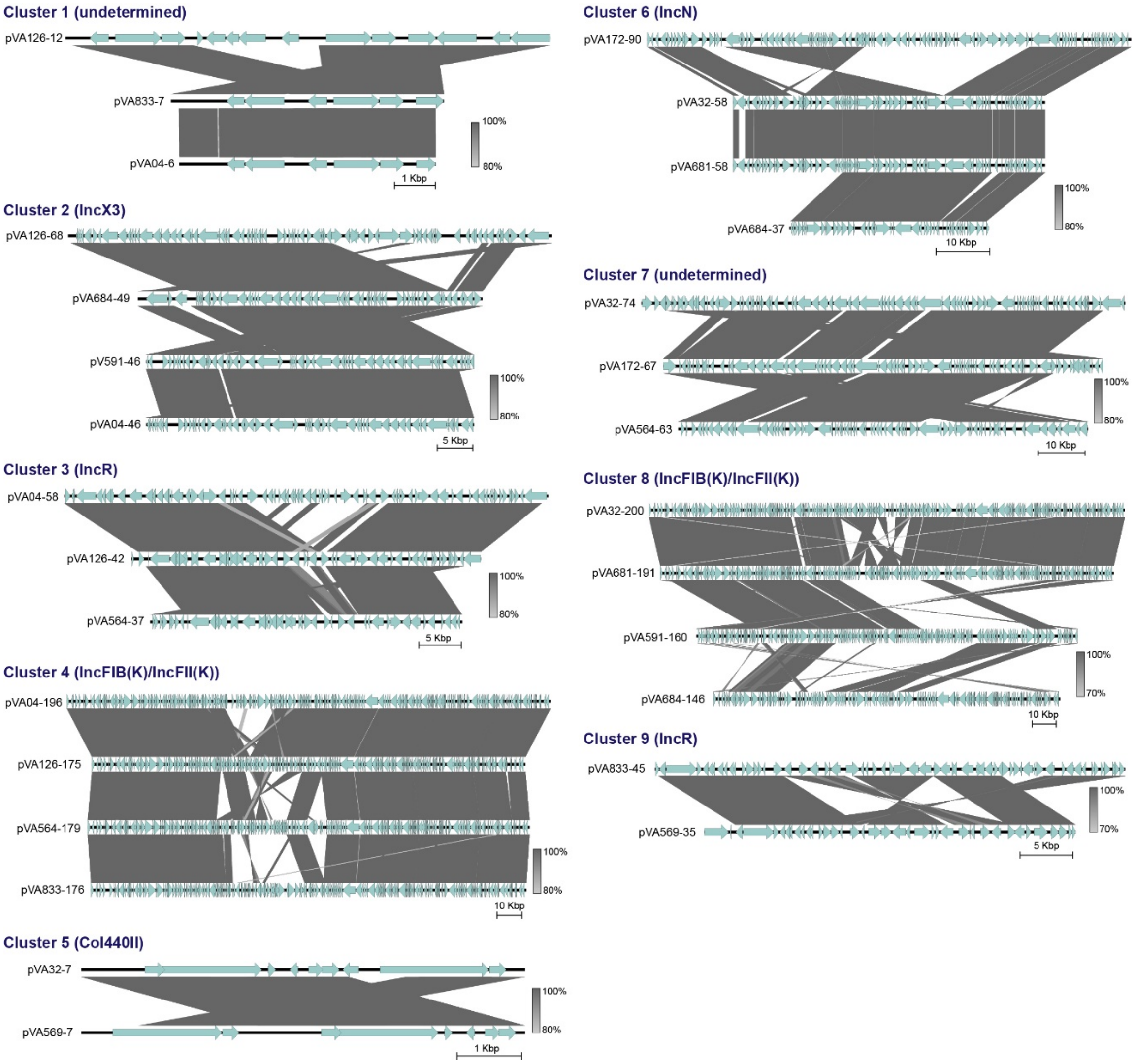


**Supplementary Figure 3.** Sequence comparison among plasmids from Chilean CR-Kp isolates composing the different clusters identified through CD-HIT analysis. The alignment plots were generated using the EasyFig tool.

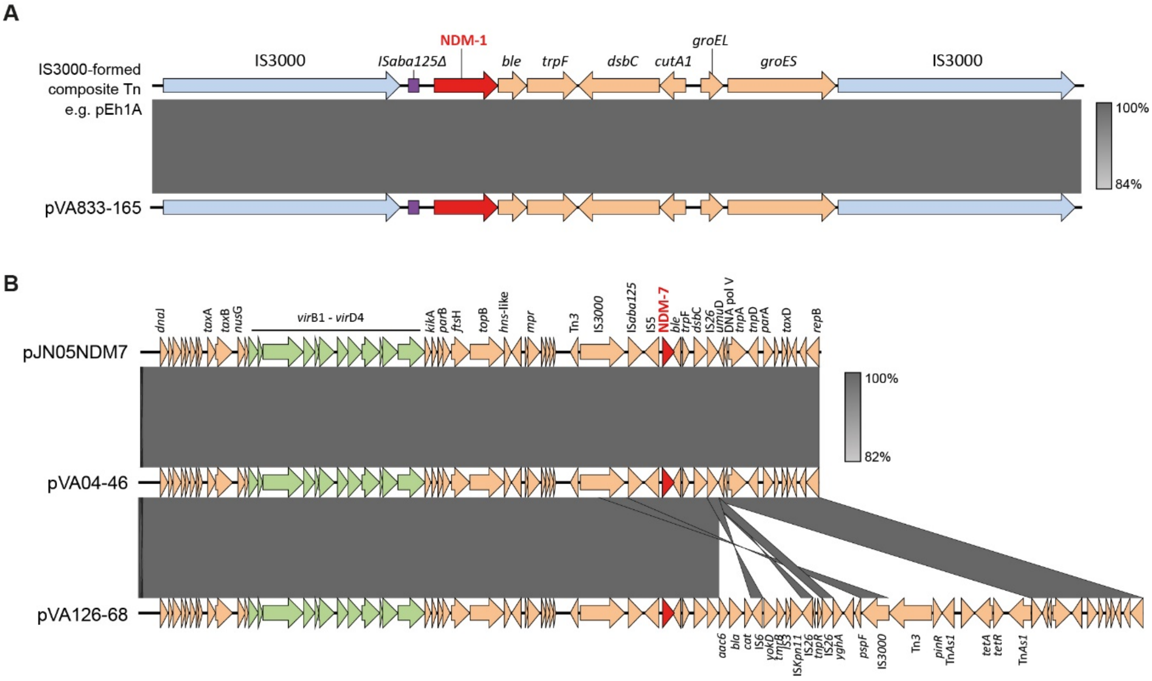


**Supplementary Figure 4.** Genetic context of the *bla*NDM-1 (A) and *bla*NDM-7 (B) genes encoded in plasmids from the Chilean CR-Kp isolates described in this study. pEh1A accession: KR822246; pJN05NDM7 accession: MH523639. The alignment plots were generated using the tool EasyFig.

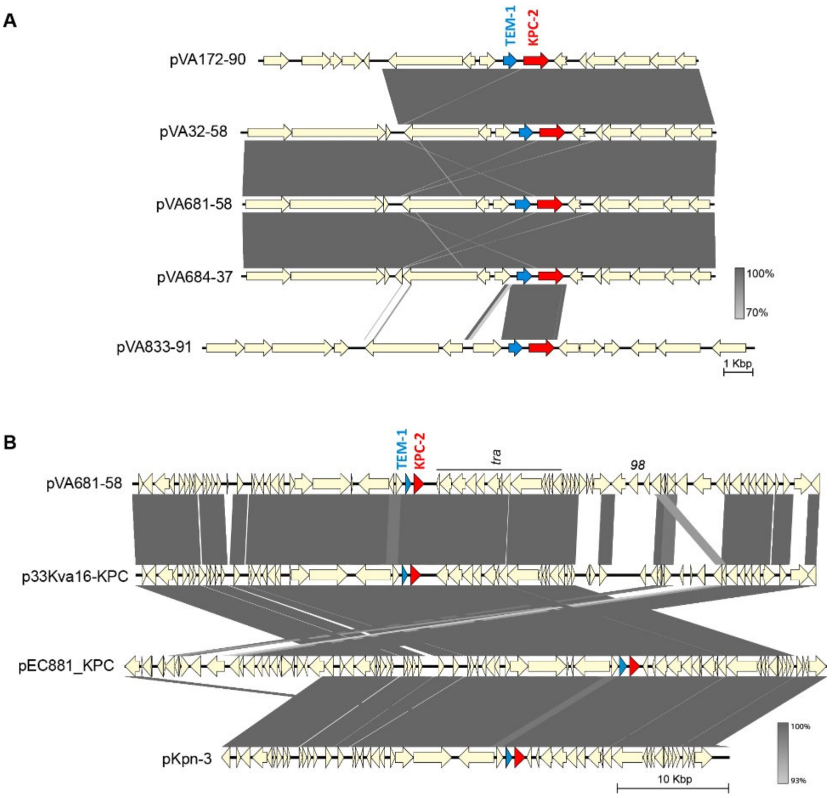


**Supplementary Figure 5.** (A) Comparison of the *bla*$_{KPC-2}$ genetic context found in different plasmids from the Chilean CR-Kp isolates described in this study. (B) Comparison of the *bla*$_{KPC-2}$ genetic context found in pVA681-58 and related plasmids described previously bearing the NTE$_{KPC}$-IIe context.

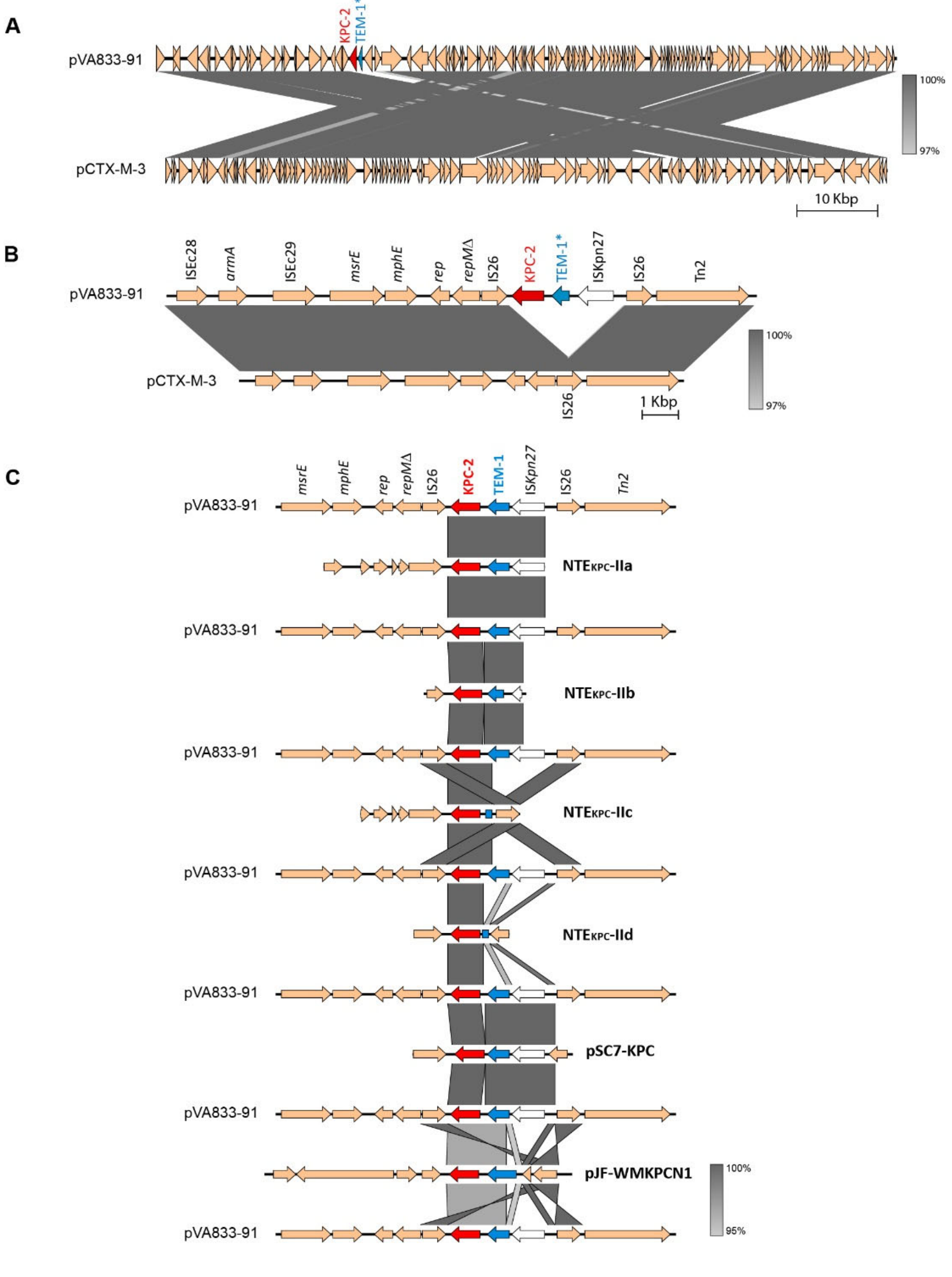


**Supplementary Figure 6.** (A) Sequence alignment between pVA833-91 and pCTX-M-3 sharing high identity. (B) Comparison of the *bla*$_{KPC-2}$ genetic context in pVA833-91 and the corresponding region in pCTX-M-3 lacking this carbapenemase gene. (C) Comparison of the *bla*$_{KPC-2}$ genetic context found in pVA833-91 and other class-II contexts (known as NTE$_{KPC}$-II) reported previously for this gene [38], indicating that this would correspond to a novel genetic environment for *bla*$_{KPC-2}$. pSC7-KPC (accession NZ_CP030267) and pJF-WMKPCN1 (accession NZ_KX881941) are plasmids recently proposed to host novel unclassified contexts.

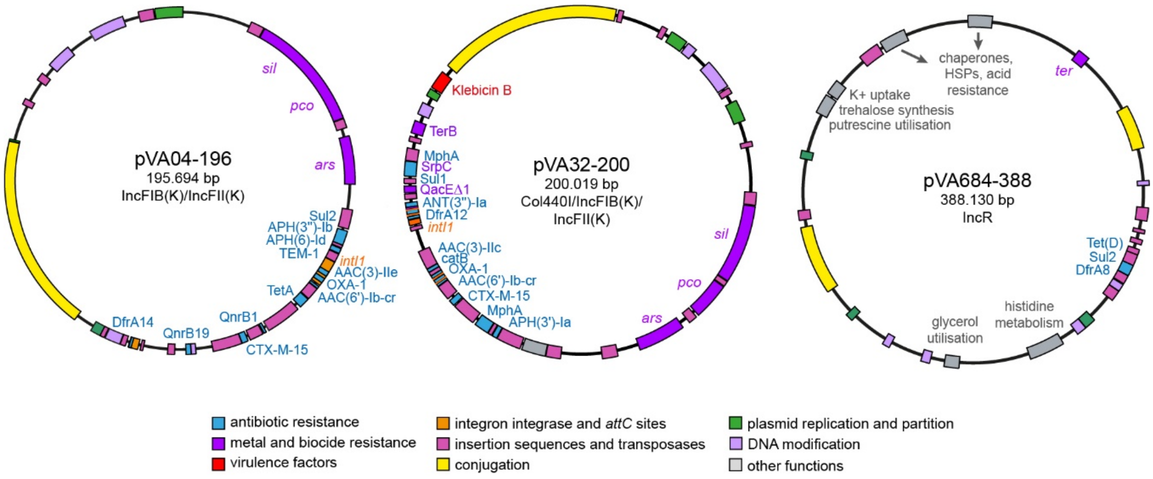


**Supplementary Figure 7.** Plasmids encoding different factors which could contribute to bacterial resistance, virulence and fitness during host infection, found in the CR-Kp isolates described in this study.
